# Concurrent encoding of sequence predictability and event-evoked prediction error in unfolding auditory patterns

**DOI:** 10.1101/2023.10.06.561171

**Authors:** Mingyue Hu, Roberta Bianco, Antonio Rodriguez Hidalgo, Maria Chait

## Abstract

Human listeners possess an innate capacity to discern patterns within rapidly evolving auditory sequences. Core questions, at the forefront of ongoing research, focus on the mechanisms through which these representations are acquired and whether the brain prioritizes or suppresses predictable sensory signals.

Previous work, using fast sequences (tone-pips presented at a rate of 20Hz), revealed sustained response effects that appear to track the dynamic predictability of the sequence. Here we extend the investigation to slower sequences (4Hz), permitting the isolation of responses to individual tones. Stimuli were 50ms tone-pips, ordered into random (RND) and regular (REG; a repeating pattern of 10 frequencies) sequences; Two timing profiles were created: in ‘fast’ sequences tone-pips were presented in direct succession (20 Hz); in ‘slow’ sequences tone-pips were separated by a 200ms silent gap (4 Hz).

Naive participants (N=22; both sexes) passively listened to these sequences, while brain responses were recorded using magnetoencephalography (MEG). Results unveiled a heightened magnitude of sustained brain responses in REG when compared to RND patterns. This manifested from three tones after the onset of the pattern repetition, even in the context of slower sequences characterized by extended pattern durations (2500ms). This observation underscores the remarkable implicit sensitivity of the auditory brain to acoustic regularities. Importantly, brain responses evoked by single tones exhibited the opposite pattern - stronger responses to tones in RND compared to REG sequences. The demonstration of simultaneous but opposing sustained and evoked response effects reveals concurrent processes that shape the representation of unfolding auditory patterns.

**Significance Statement:** Humans excel at detecting predictable patterns within sound sequences, a process crucial for listening, language processing, and music appreciation. However, questions persist about the underlying neural mechanisms and the specific information monitored by the brain.

Our study addresses these questions by analysing magnetoencephalography (MEG) signals from participants exposed to predictable and unpredictable tone-pip patterns. We found that the MEG signal simultaneously captures two crucial aspects of predictability tracking.

Firstly, sustained MEG activity, tracking the sequence’s evolution, dynamically assesses pattern predictability, shedding light on how the brain evaluates reliability. Secondly, phasic MEG activity, reflecting responses to individual events, shows reduced activity to predictable tones, aligning with the idea that the brain efficiently encodes and anticipates upcoming events in predictable contexts.

## Introduction

The physical rules that govern the environment and impose constraints on its agents result in statistically structured, predictable sensory signals. The brain is hypothesized to have developed the capacity to rapidly detect and track the regularities within these signals (de Lange et al., 2018; Press et al., 2020). This ability plays a crucial role in the comprehension of our surroundings, facilitating efficient recognition and processing of incoming information, to empower us to respond rapidly and adaptively to changing circumstances.

The auditory system, in particular, has demonstrated remarkable tuning to regularities across various time scales and dimensions (Bendixen, 2014; Heilbron & Chait, 2018; Carbajal & Malmierca, 2018; Asokan et al, 2019; Fitzgerald & Todd, 2020).This plays a crucial role in our ability to understand spoken language (Arnal and Giraud, 2012), appreciate the nuances of musical compositions (Koelsch et al., 2019) and make sense of the complex soundscape that surrounds us. However, core questions regarding the mechanisms through which regularity is discovered and tracked remain elusive. In particular, pivotal issues revolve around whether the brain chooses to prioritize or suppress predictable sensory signals (Press et al, 2020).

Barascud et al. (2016); see also (Sohoglu and Chait, 2016; Southwell et al., 2017; Herrmann and Johnsrude, 2018; Herrmann et al., 2019) provided insight into the brain’s automatic ability to detect the emergence of predictable acoustic structure by examining low-frequency activity in the M/EEG signal. Using rapidly unfolding (20 Hz) tone-pip sequences that contained transitions from a random (RND) to a regularly repeating pattern (REG), they observed that a gradual increase in sustained power accompanies the emergence of repeating structures. The timing of the differentiation between REG and RND sequences (3 tones after the first cycle) was consistent with that predicted by an ideal observer model (Pearce, 2005; Harrison et al., 2020), demonstrating statistically efficient processing of structure even when not behaviourally relevant (Barascud et al., 2016).

The sustained response effect is interesting for several reasons: Firstly, it suggests that the brain encodes the inherent state of the stimulus (RND vs REG) rather than merely registering changes in the environment. Secondly, the observed ***increase*** in sustained power during structure discovery challenges our understanding of how the brain processes and represents predictability. Specifically, it appears to contradict expectations derived from predictive coding frameworks (e.g. Friston, 2005, 2009; Rao & Ballard, 1999), where predictable information is typically associated with ***reduced*** neural activity, as the brain can efficiently encode and predict upcoming events (de Lange et al., 2018). Barascud et al. showed that the sustained response, underpinned by activation in the auditory cortex, hippocampus, and inferior frontal gyrus, increases with the predictability of the ongoing stimulus sequence. This prompted the hypothesis that it might reflect the process of tracking the inferred reliability of the unfolding input (‘precision’; the accuracy, or conversely the ‘expected uncertainty’ with which future inputs can be predicted, O’Reilly et al., 2013) whereby predictable sensory streams are associated with heightened sensitivity.

Several issues need to be addressed for a better interpretation of the sustained response. Firstly, it is important to consider that the effects observed may be specific to the rapid sequences used in Barascud et al. (2016). Other research (e.g. reviewed by (de Lange et al., 2018; Heilbron and Chait, 2018) has focused on slower patterns, which may elicit different neural responses. Secondly, it is crucial to determine whether the observed effect primarily reflects a shift in background neural activity or if it also extends to modulations of responses to individual events due to their integration within the structured sequence.

To address these questions, the current study expands upon the original stimulus by introducing silent gaps between successive tones (Figure 1, 2). We aim to explore the generality of the sustained-response effects across different temporal scales and provide a clearer understanding of the mechanisms involved in the processing of structured auditory sequences.

**Figure 1.**
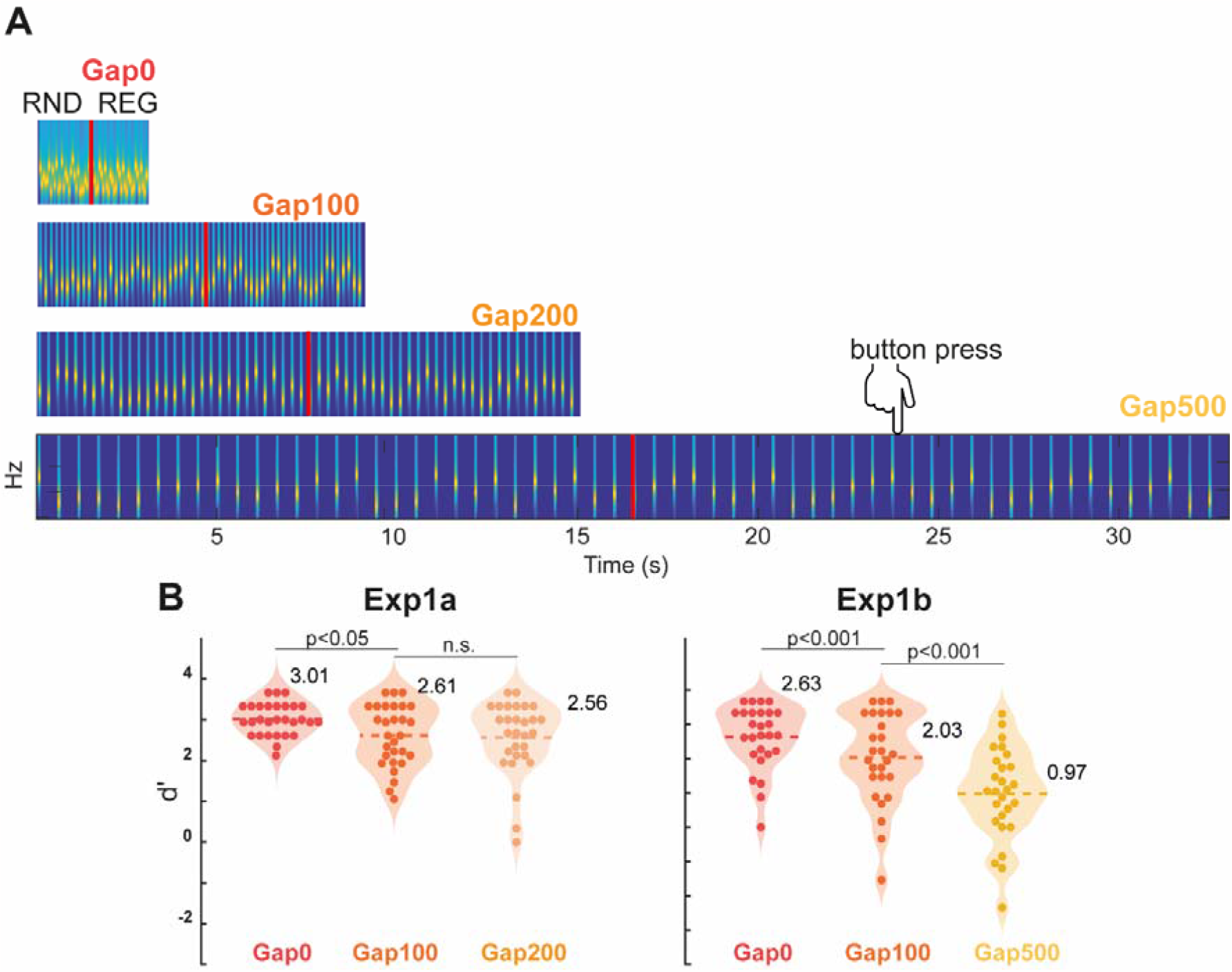
Behavioural experiment. [A] Examples of the four gap duration stimuli (to scale). RNDREG sequences are plotted (the stimulus set also contained 50% no-change RND sequences). Four gap duration conditions are used (0, 100, 200 and 500 ms), resulting in regularity cycles of 500, 1500, 2500 and 5500 ms, respectively. Participants listened to the sound sequences and were instructed to press a keyboard button as soon as they detected the emergence of a REG pattern; indicated with a red line. [B] Behavioural performance. Performance steadily declined with increasing gap duration. Generally good performance (mean d’>2) was seen for the Gap200 condition and it was therefore chosen for the MEG experiment.

**Figure 2.**
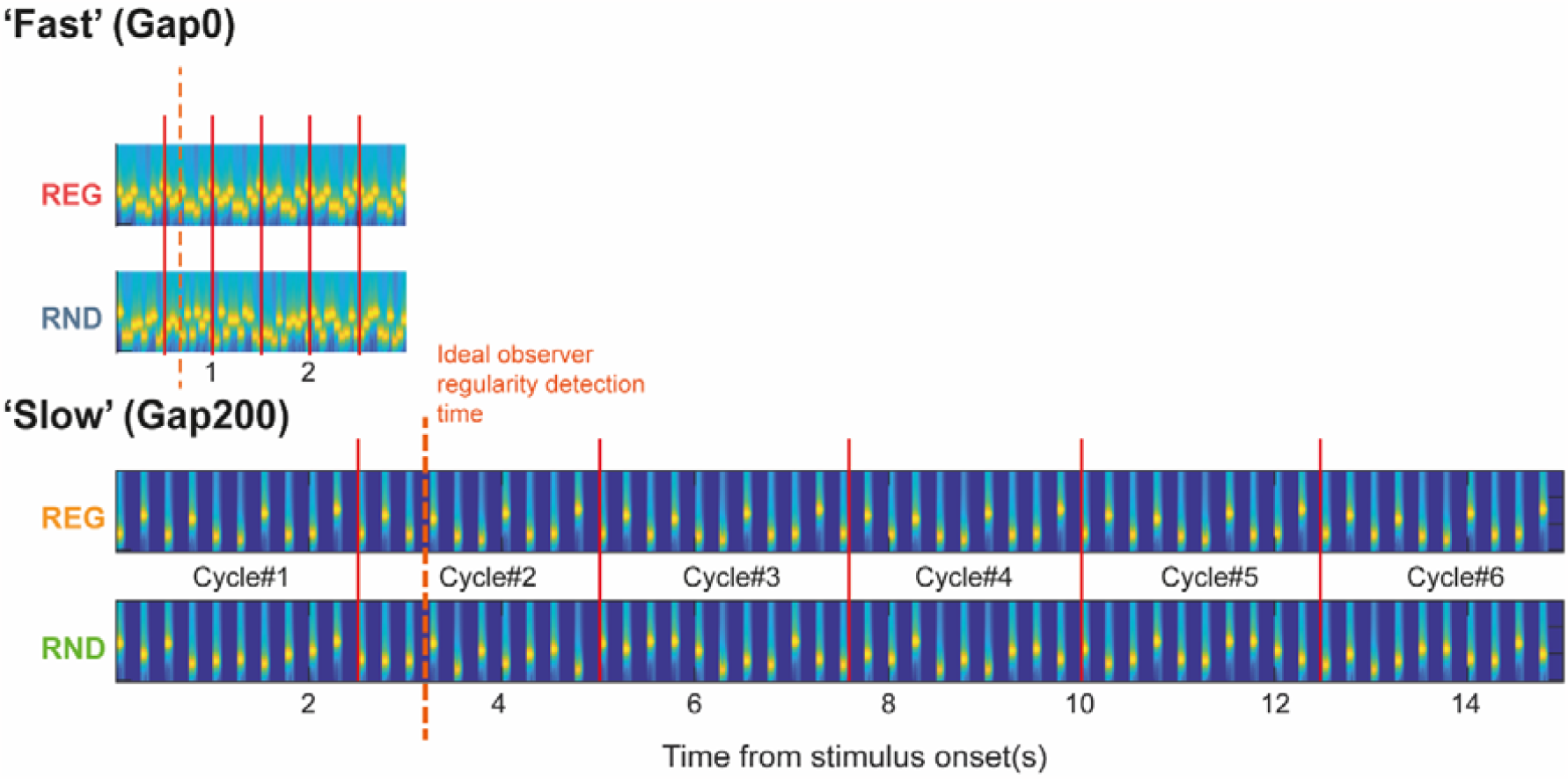
Examples of stimuli in the MEG experiment (to scale). All stimuli consisted of 60 tones (6 regularity cycles in REG sequences; red lines). ‘fast’ sequences were 3 s long; ‘slow’ sequences were 15 s long. Naive participants listened to the sound sequences passively and were instructed to focus on a visual task. If brain responses track the emergence of regularity, responses REG and RND sequences should be differentiated following cycle#1. Ideal observer REG detection latency (∼3 tones into the 2^nd^ cycle, e.g. Barascud et al, 2016) is indicated with a dashed line.

## Methods

### Experiment 1 - Online behavioural study

The behavioural study was designed to probe how the introduction of silent gaps between tones affects explicit pattern detection. We sought to pinpoint an optimal gap duration that is sufficiently long to allow us to isolate responses to individual tones, yet brief enough to maintain high-performance levels in pattern detection.

### Stimuli

Stimuli were sequences of 50-ms tone-pips (gated on and off with 5-ms raised cosine ramps) drawn from a pool of 20 values equally spaced on a logarithmic scale between 222 and 2000 Hz (12% steps). The order in which these tone-pips were successively distributed defined two different sequence types. **RND** sequences consisted of 20 tone-pips (sampled from the full pool) arranged in random order. Each tone-pip occurred equi-probably across the sequence duration. **RNDREG** sequences contained a transition between a RND sequence, and a regularly repeating pattern (REG). REG consisted of 10 different tone-pips, randomly chosen from the full pool on each trial, and repeated in 3 identical cycles. The RND to REG transition always occurred after 30 tone-pips. Opting for this method, as opposed to a variable transition time, ensured a consistent context (in terms of frequency information available) both preceding each transition and across different gap duration conditions. RND and RNDREG sequences were generated anew for each trial and presented equi-probably throughout the experiment. Therefore, the occurrence of a transition in any given trial was unpredictable. The amplitude of each tone pip was normalized to yield an approximately similar perceived loudness (Moore, 2014). Across blocks, the inter-tone-intervals were manipulated to form four conditions (Figure 1A): **Gap0** (continuous presentation), **Gap100** (a 100 ms gap inserted between tones), **Gap200** (a 200 ms gap inserted between tones), **Gap500** (a 500 ms gap inserted between tones).

Two control stimuli were also included: sequences of contiguous (no silent gap) tone-pips of a fixed frequency (**CONT**) that lasted 4000 ms, and sequences with a step change in frequency partway through the trial (**STEP**: the change always occurred after 2000 ms). These were used to measure individuals’ response time to simple acoustic changes and served as ‘catch trials’ to assess task engagement.

### Procedure

The experiment was implemented online using the Gorilla Experiment Builder (www.gorilla.sc). Before the main task, participants completed a headphone screening task (Milne et al., 2020) to ensure they were using appropriate audio equipment. They then received an explanation of the task and completed a practice session. Due to length constraints, the experiment was divided into two parts, performed by two different groups of participants. Experiment 1a contained the Gap0, Gap100 and Gap200 conditions along with the control stimuli (STEP and CONT; see above). Experiment 1b contained the Gap0, Gap100 and Gap500 conditions, along with the control stimuli.

Participants were instructed to respond, by pressing a keyboard button, as soon as possible once they had detected a RNDREG transition or a STEP. To motivate participants to focus on the task, they were given feedback on their accuracy and speed after each trial. A small monetary bonus was given for each correct response (Bianco et al., 2021).

In each experiment, three blocks of 40 trials were delivered. Each block contained the following sequence types: 15 RNDREG, 15 RND, 5 STEP, and 5 CONT. The first block always presented the Gap0 condition. This block lasted 5 minutes. Thereafter, listeners completed the other two blocks (Gap100 and Gap200 in experiment1a, Gap100 and Gap500 in experiment1b) in random order. Starting with Gap0 ensured that all participants experienced the easiest condition first and had adequate opportunity to practice the regularity detection task, reducing the likelihood of frustration and dropout that may occur if participants are immediately faced with the most difficult condition. The main task in experiment 1a lasted about 20 minutes, and that in experiment 1b lasted about 30 minutes.

### Participant Rejection Criteria

Previous work (Barascud et al., 2016; Bianco et al., 2020) demonstrated that participants are sensitive to the emergence of regularity in RNDREG sequences, exhibiting high sensitivity and rapid detection time (usually responding within two regularity cycles). Due to the online nature of the present experiments and associated reduced control over participants’ environments, equipment, and engagement (Bianco et al., 2021), it was important to implement a series of rejection criteria to make sure that data reflect true sequence tracking sensitivity. Therefore, subjects were excluded from the experiment following the below criteria:

1. Failure on the Headphone screen: We used the task introduced by Milne et al. (2020). Participants who failed the test did not proceed to the main experiment.
2. Low performance in the practice run: To ensure participants understand the task, 24 trials with no gap (10 RNDREG, 10 RND, 2 CONT and 2 STEP) were given. Participants did not proceed to the main task if their correct response rate was below 80% in the practice task. This ensured that those participants who proceeded to the main experiment could detect the REG transitions, thus allowing us to focus on how performance is affected by increasing the gaps between tones. Our previous experience with similar stimuli in lab settings (see e.g. Barascud et al, 2016; Bianco et al, 2020) suggests that the vast majority of young participants can achieve ceiling performance. We, therefore, reasoned that those online participants who perform below 80% are likely not sufficiently engaged with the task (i.e. distracted, not following instructions, etc).
3. Of those participants who completed the full experiment, we rejected subjects who failed to respond to STEP trials (allowing at most one miss per block) or whose RT to STEP trials fell above 2 STDEV relative to the group mean. Failure to respond quickly to the (easy) STEP trials indicated low task engagement.

### Participants

Two participant groups were recruited via the Prolific platform (https://www.prolific.co/). 29 subjects were included in the analysis of experiment1a (7 females; average age, 24.3 ± 4.79 years). 27 subjects were included in the analysis of experiment1b (6 females; average age: 22 ± 4.69 years).

### Experiment 2 - MEG in naïve passively listening participants

#### Stimuli

Stimuli (Figure 2) were generated similarly to those in experiment 1. To reduce the duration of the (passive listening) MEG experiment, we focused on REG and RND sequences, without transitions. Sensitivity to regularity is investigated by comparing brain responses to the onset of REG and RND sequences. During the initial portion of the sequence (first cycle in REG), responses to the two sequence types should be identical, with differences emerging as soon as the auditory system has discovered that the pattern is repeating. Ideal observer modelling (Barascud et al., 2016; Harrison et al., 2020) suggests that about 3 tones, following the first cycle, are needed for the transition to be statistically detectable. REG sequences were generated by randomly selecting (without replacement) 10 frequencies from the pool and iterating that order to create a regularly repeating pattern. RND sequences consisted of a random succession of 10 tones, newly selected on each trial. All stimuli contained 60 tone-pips. Two timing conditions were used: in ***‘fast’*** sequences tone-pips were presented in direct succession (20 Hz rate; 500ms REG cycle duration; 3 s overall sequence duration); in *‘slow’* sequences tone-pips were separated by a 200 ms silent gap (4 Hz rate; 2500ms REG cycle duration; 15 s overall sequence duration). One hundred instances of each condition were presented. Sequences were generated anew for each trial such that each stimulus was created of the same frequency “building blocks” (random selection of 10 out of 20 frequencies). Condition presentation was fully randomized.

### Procedure

The experiment was controlled with the Psychophysics Toolbox extension in MATLAB (Kleiner et al., 2007). All auditory stimuli were presented binaurally via tube earphones (EARTONE 3A 10 Ω; Etymotic Research) inserted into the ear canal, with the volume set at a comfortable listening level, adjusted for each participant.

The experiment lasted 40 minutes. Participants listened passively to the stimuli (presented in random order with an ISI jittered between 1400-1800 ms) and engaged in an incidental visual task. The task consisted of landscape images, grouped in triplets (the duration of each image was 5 s, with 2 s ISI between trials during which the screen was blank). Participants were instructed to fixate on a cross in the centre of the screen and press a button whenever the third image was identical to the first image (10% trials). The visual task served as a decoy task for diverting subjects’ attention away from the auditory stimuli. Participants were naïve to the nature of the auditory stimuli and encouraged to focus on the visual task. Feedback was displayed at the end of each block. The experimental session was divided into six 12 min blocks. Participants were allowed a short break between blocks but were required to remain still.

### Participants

23 naïve subjects participated in the study. One of them was discarded due to excessive noise in the data. Data from 22 participants (11 females; average age, 25.14 ± 4.61 years) are reported below.

### Data recording and pre-processing

Magnetic signals were recorded using CTF-275 MEG system (axial gradiometers, 274 channels; 30 reference channels; VSM MedTech). The acquisition was continuous, with a sampling rate of 600 Hz. Offline low-pass filtering was applied at 30 Hz (all filtering in this study was performed with a two-pass, Butterworth filter with zero phase shift). All pre-processing and time domain analyses were performed using the fieldtrip toolbox (Oostenveld et al., 2011). To analyse time domain data, we selected the **40 most responsive channels** for each subject. This was done by collapsing across all conditions and identifying the M100 component of the onset response (Näätänen and Picton, 1987; Stufflebeam et al., 1998; Näätänen et al., 2011; Gorina-Careta et al., 2021), as a source-sink pair located over the temporal region of each hemisphere. For each subject, the 40 most strongly activated channels at the peak of the M100 (20 in each hemisphere; 10 in each sink/source) were considered to best reflect auditory activity and thus selected for all subsequent time-domain analyses. This procedure served the dual purpose of enhancing the relevant response components and compensating for any channel misalignment between subjects.

We report two time-domain analysis pipelines:

### Whole sequence analysis

Initially, we focused on responses to the entire sequence. Low-frequency activity is of prime importance as a possible marker of predictability tracking (Barascud et al., 2016; Southwell et al., 2017). Therefore, no high-pass filter was used. Data were segmented into epochs from 200ms before onset to 1000ms post offset (yielding epochs of 4200ms and 16200ms in ‘*fast’* and ‘*slow’* conditions, respectively). Epochs containing artefacts were removed (based on variance summary statistics) variance using Fieldtrip’s manual visual artefact rejection function. Around 5% of epochs were removed from each subject (range 0-10%). The remaining epochs were then averaged by condition. To help denoise the data from *‘slow’* conditions (low-frequency drift artefacts) denoising source separation (DSS) analysis was applied to maximize reproducibility across trials. (Särelä and Valpola, 2005; de Cheveigné and Simon, 2008; de Cheveigné and Parra, 2014). For each subject, the three most significant components (i.e., the three ‘most reproducible’ components across trials) were kept and projected back into sensor space.

### The single-tone response analysis

A subsequent analysis focused on responses to individual tones in REG vs. RND sequences in the *‘slow’* sequences. To identify activity associated with individual tone-evoked responses which might be masked by the sustained activity, the raw data were high-pass filtered at 2Hz. Filtered data were then cut into individual tone epochs, from 50 ms before the onset of the tone, to 200 ms post onset. Responses from tones within each cycle were averaged, yielding 6 time series per condition per subject (tones in Cycle#1, Cycle#2, etc.). Time series were baselined based on pre-tone onset activity.

### Statistical analysis

The time domain data are summarised as root-mean square (RMS) across the 40 selected channels for each subject (see above). The RMS is a useful summary signal, reflecting the instantaneous power of the neural response irrespective of its polarity. Group RMS (RMS of individual subject RMSs) is plotted; statistical analysis was always performed across subjects.

To evaluate differences between conditions (RND vs REG), the RMS differences at each time point were computed for each participant, and a bootstrap re-sampling (Efron and Tibshirani, 1998) was applied (10000 iterations) on the entire epoch. Significance was inferred by inspecting the proportion of bootstrap iterations that fell above or below zero, here p=0.01 was used as a threshold.

#### Source analysis

To estimate the brain sources that underly the observed time domain effects at the sensor level, we performed source reconstruction using the standard approach implemented in SPM12 (Litvak and Friston, 2008; López et al., 2014; Bartha-Doering et al., 2015) Sensor-level data were converted from Fieldtrip to SPM. By using 3 fiducial marker locations, the data were co-registered to a generic 8196-vertex inverse-normalised canonical mesh warped to match the SPM’s template head model based on the MNI brain (Ashburner and Friston, 2005). This had the advantage of providing a one-to-one mapping between the individual’s source-space and the template space, facilitating group analyses (Litvak and Friston, 2008). The forward model was solved with a single shell forward head model for all subjects. Source reconstruction was performed using the multiple sparse priors (MSP) model that assumes that activity can be expressed in multiple patches or covariance components, each of which has an associated hyperparameter (Litvak and Friston, 2008; López et al., 2014; Bartha-Doering et al., 2015). These were optimised with greedy search (GS) technique (Litvak and Friston, 2008) by iterating over successive partitions of multiple sparse priors to find the set yielding the best fit (here we specify a total of 512 total dipoles). The MSP model was used to identify distributed sources of brain activity, hence the 2 conditions (RND and REG) were inverted together.

We were interested in capturing the sources underlying two aspects of the data:

1. The discovery of regularity (REG vs RND) in the *‘fast’* sequence evoked response. The analysis used DSSed data (de Cheveigné and Parra, 2014), with the three most reproducible components projected back into sensor space and used for the inversion. Trials were averaged by condition and the inverse estimates were obtained for the two conditions together using an interval of 300ms between 665 and 965 ms post-stimulus onset. The interval was chosen to coincide with the timing of divergence between the REG and RND conditions as seen in the time domain analysis (Figure 3).

**Figure. 3.**
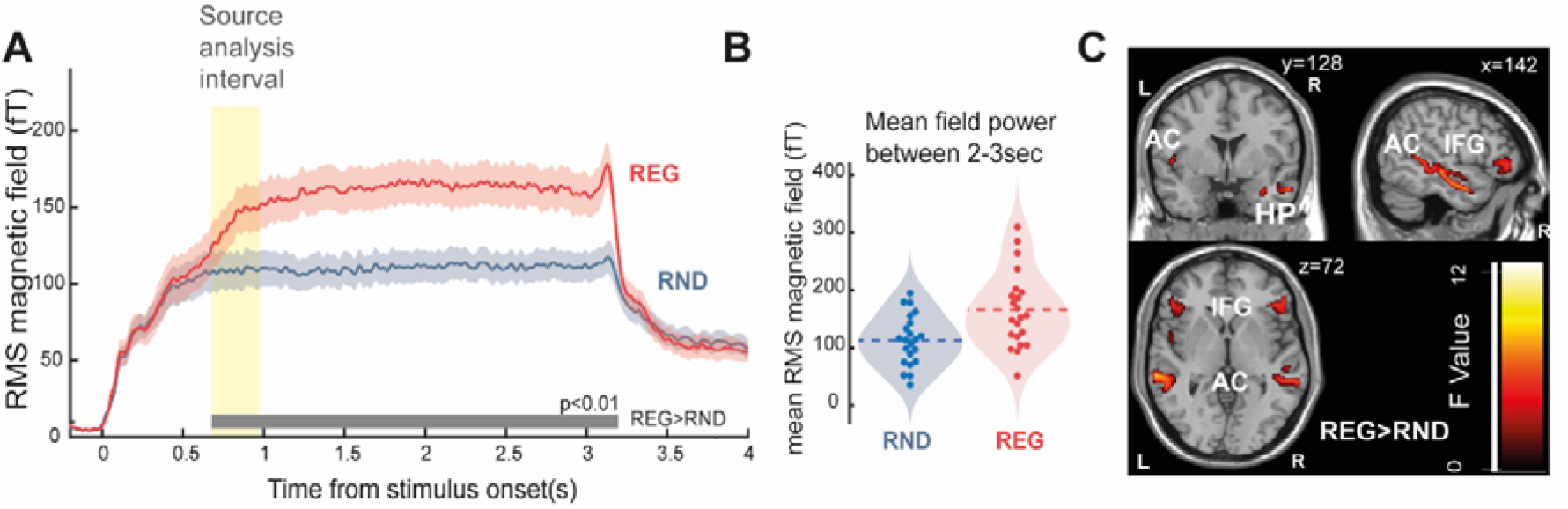
MEG response to ‘fast’ (Gap0) sequences. [A] The full stimulus epoch, from stimulus onset (t = 0s) to offset (t = 3s). The shaded area around the traces indicates the standard error of the mean. The grey horizontal line indicates time intervals where a significant difference is observed between the two conditions (p<0.01). Yellow highlighting indicates the interval (665 ms to 965 ms) used for source analysis in [C]. [B] Mean sustained response power computed during the last second of stimulus presentation (2-3 s post-onset) and averaged over trials for each subject in RND and REG conditions. [C] Source analysis. Group F map for the REG > RND during the rising slope of the sustained response (yellow shaded area in A), thresholded at p = 0.05 (uncorrected).
2. The effect of regularity (REG vs. RND) on the individual tone responses in *‘slow’* sequences. A similar analysis pipeline as that described above was used. This analysis focused on the interval between 5 and 15 s – from the 3^rd^ cycle of the REG until offset, i.e., where the regularity in REG stimuli was well established (theoretically, and, as seen in the time domain data, regularity is discovered partway through the 2^nd^ cycle and well established by the 3^rd^ cycle). The filtered raw signal (2-30 Hz), epoched over 0-200ms post tone onset and averaged across tone presentations, was used for the inversion. The interval was chosen to coincide with the largest possible time window post tone onset to allow the algorithm to encompass all brain sources responsible for generating the response (Henson et al., 2011).

After inversion, source activity for each condition was projected to a three-dimensional source space and smoothed [12-mm full width at half maximum (FWHM) Gaussian smoothing kernel] to create Neuroimaging Informatics Technology Initiative (NIfTI) images of source activity for each subject. At the second level of statistical analysis, the two conditions (REG vs RND) were modelled with the within-subject factor Regularity (REG / RND). Statistical maps of the contrast were thresholded at a level of p < 0.05 uncorrected (F contrasts) across the whole-brain volume. Relevant brain regions were identified using the AAL3 toolbox (https://www.oxcns.org/aal3.html).

## Results

### Behavioural performance reveals good sensitivity to regularity even following the introduction of silent gaps between tones

We tested how pattern detection ability is affected by the introduction of a silent gap of increasing length between successive tone pips. Figure 1B shows performance (quantified as d’ sensitivity score) for each condition in experiments 1a and 1b. With increasing gap duration, an overall gradual worsening of performance was observed. A repeated measures ANOVA over the three gap duration conditions in experiment 1a confirmed a main effect of condition [F (2, 56) = 3.814, η2 = .123, p = .026]. Post hoc tests (Bonferroni corrected) indicated a significant difference between Gap0 and Gap100 conditions [p = .034] and between Gap0 and Gap200 conditions [p = .026]. No difference between Gap100 and Gap200 was seen [p = 1]. In general, most participants achieved a d’ above 2 in the Gap200 condition, revealing a largely conserved sensitivity even though the duration of the pattern increased five-fold from 500ms in Gap0 to 2500ms in Gap200. In experiment 1b we further tested the performance for silence gaps of 500 ms. A repeated measures ANOVA with factor Gap (0, 100, 500 ms) confirmed a main effect of condition [F(2, 52) = 33.687, η2 = .564, p < .001]. Post hoc (Bonferroni corrected) comparisons indicated significantly worse performance in Gap100 [p=.025] and Gap500 [p<.001] compared to Gap0, and between Gap100 and Gap500 [p < .001].

Overall, the pattern of results is consistent with a slow decline in performance for gaps up to 200ms and a steeper drop thereafter. We, therefore, selected the 200ms gap duration for the MEG experiments (in naïve distracted listeners) below.

### The emergence of regularity is associated with an increase in sustained MEG activity

The Group RMS (mean of all subjects’ RMSs) of the evoked response to the ‘*fast’* sequences are shown in Figure 3A. The brain response presents prototypical onset activity, followed by a subsequent rise to a sustained response that persists until offset. A pronounced offset response is seen about 100 ms after sound cessation. Fluctuations at 20 Hz, reflecting the tone presentation rate, are visible in the sustained portion of the response. In line with previous observations (Barascud et al., 2016; Southwell et al., 2017; Southwell and Chait, 2018), REG shows an increased sustained response when compared with RND. The timing at which the response to REG diverges from RND is considered to reflect the information required to detect the regularity. A significant difference between conditions emerged after 665 ms, (13 tone-pips, 1.3 cycles). This estimate is consistent with previous modelling work (Barascud et al., 2016; Harrison et al., 2020) which demonstrated that an ideal observer model required 3-4 tones following the first cycle to detect the emergence of regularity.

Figure 3B displays the source analysis, applied over a 300 ms interval over which the REG and RND conditions begin to diverge (yellow shading in Figure 3A). The activation map (F contrast, REG>RND) demonstrates increased activity in auditory cortex (AC; bilaterally), inferior frontal gyrus (IFG; bilaterally) and hippocampus (HP; RH only). No areas were identified by using the opposite (RND > REG) contrast. Overall, the source data are largely consistent with what was previously shown by Barascud et al. 2016 for similar stimuli, confirming a distributed network spanning auditory, frontal and hippocampal sources which underlies sensitivity to regular patterns.

Responses to the ‘*slow’* (Gap200) sequences are shown in Figure 4A. Pronounced fluctuations at 4 Hz, reflecting the tone presentation rate, are clearly visible on top of the sustained response. Similar to what was observed for the ‘*fast’* sequences, a difference in sustained response emerges between REG and RND when the REG pattern begins to repeat (after 2500ms). This effect is much smaller, however. To separate the sustained response from phasic activity associated with tone-evoked responses, the data were low pass filtered (0-2Hz; Figure 4B). A significant difference between conditions emerged after 13 tones (3266 ms) consistent with the observations from the *‘fast’* sequence above. This suggests that irrespective of the rate at which tones are presented (at least within the range tested here), regularity detection requires a constant amount of information (as measured in number of tones pips). However, it is notable that the sustained difference between REG and RND conditions in the ‘*slow*’ sequences is smaller and rather noisier (e.g. as reflected by the discontinuous significance, see Figure 4) than in the ‘*fast*’ sequences.

**Figure. 4.**
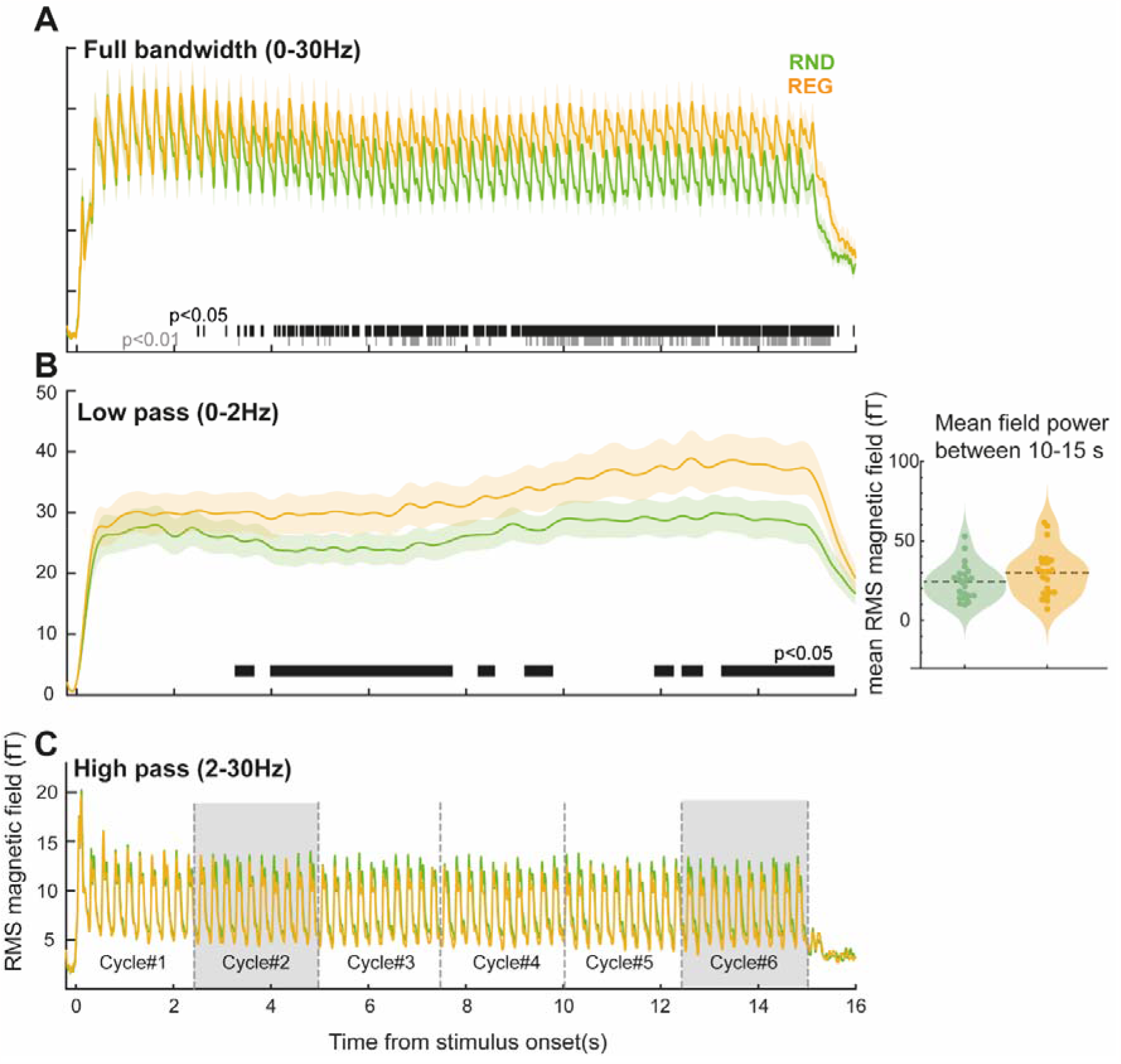
MEG response to ‘slow’ (Gap200) sequences. [A] Wideband; 0-30Hz. The entire stimulus epoch (16s) is plotted. A sustained difference between responses to REG and RND sequences emerges from ∼ 3s post-onset. Responses evoked by individual tones (4Hz) are observed throughout the epoch. [B] Low pass filtered responses (0-2Hz) focusing on the slow sustained response activity. The horizontal black and grey lines denote time intervals where a significant difference is observed between conditions (p < .05 and p<.01, respectively). Mean sustained response power computed between 10-15 s (from the 5th cycle onwards) post-onset for each individual in each condition is shown on the right. [C] High pass filtered activity, with clearly visible responses to individual tones. The 6 REG cycles analysed in Figure 5 are indicated. Shaded areas are those plotted in Figure 5B, C.

Overall, the MEG results demonstrate that passively elicited brain responses to REG relative to RND sequences are associated with significantly stronger sustained response magnitude, including when pattern durations are long (2500 ms in *‘slow’* sequences).

### Responses to individual tones are decreased in REG relative to RND sequences

To focus on phasic activity associated with responses to individual tones, sequence-evoked responses were high pass filtered at 2Hz (Figure 4C) and tone-centred epochs were extracted (from 50ms pre-tone-onset to 200ms post-tone-onset). The main analysis (Figure 5), focused on tones presented in each cycle of the REG sequences (see indicated in Figure 4C; 0-2.5 s; 2.5-5 s; 5-7.5 s; 7.5-10 s; 10-12.5 s; 2.5-15 s), and corresponding tones in RND sequences. As expected, no differences between conditions are seen in the first cycle (cycle#1) (Figure 5A). In contrast, clear differences between tones presented in REG vs RND contexts are seen in cycle #2 onwards (Figure 5B; cycle #6 also plotted; Figure 5C). Critically, REG tones evoke *reduced* responses relative to RND tones. This effect appears to be specific to the latter part of the tone-evoked response: from ∼100ms post tone onset, i.e., during the tone-evoked M100 peak.

**Figure. 5.**
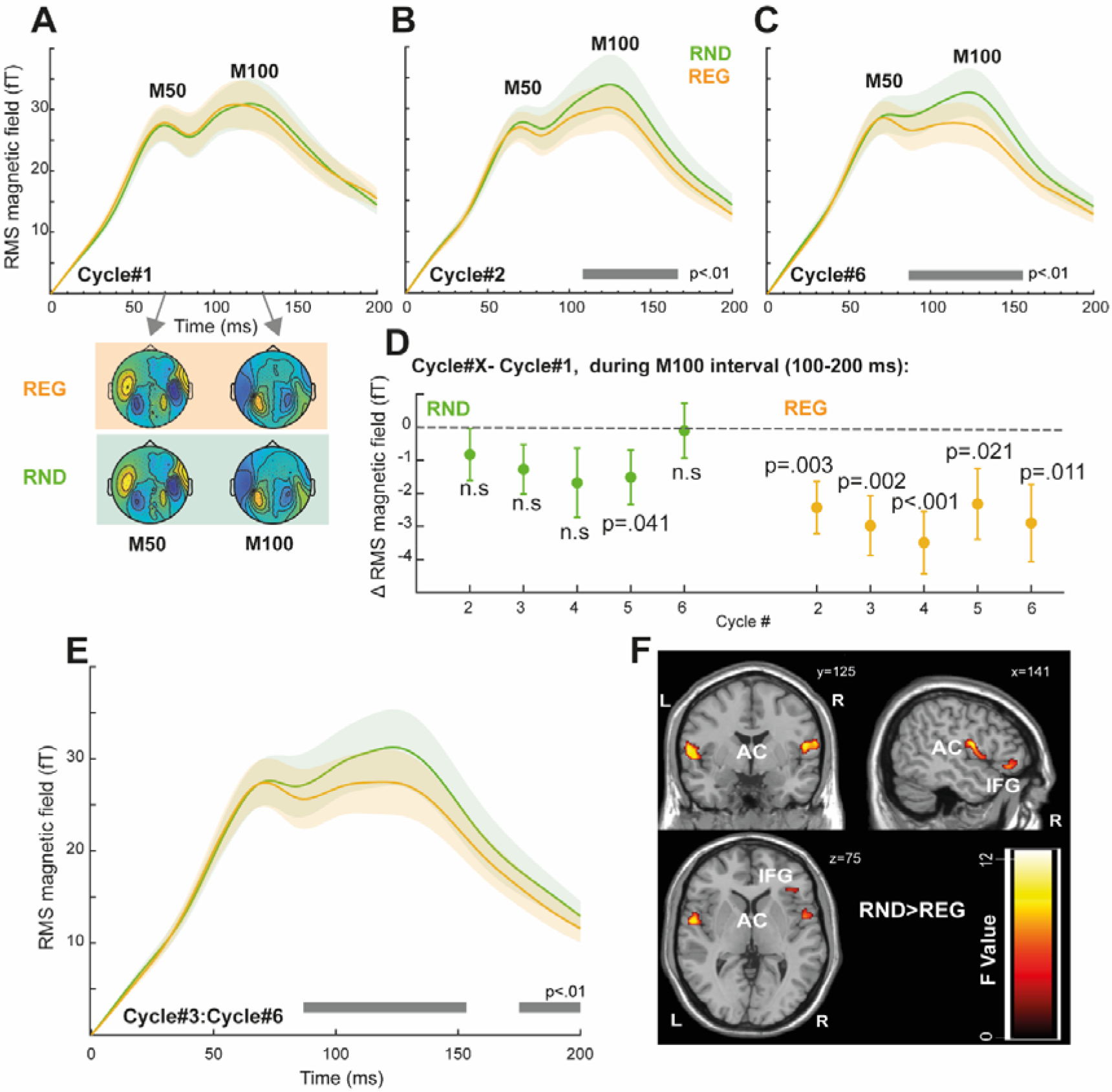
Tone evoked responses. [A] Tone-evoked responses averaged over the first 10 tones (0-2.5 s; first cycle) in the RND and REG conditions. Shading around the traces indicates the standard error of the mean. Field maps corresponding to the M50 (60-80 ms) and M100 (130-150 ms) responses are shown below. As expected, no differences are seen because the REG pattern can only be distinguished from RND following the first cycle (once the pattern starts repeating) [B] Tone-evoked responses averaged over tones presented between 2.5 - 5 s in the RND and REG conditions (‘Cycle#2). The horizontal grey line indicates time intervals where a significant difference is observed between conditions (p<.01) [C] Tone-evoked responses averaged over tones presented between 12.5 - 15 s in the RND and REG conditions (‘Cycle#6). [D] Difference from 1^st^ cycle computed (over the M100 time interval; 100-200ms) for each subsequent cycle in REG and RND. Tones presented in REG contexts show consistently reduced activity relative to the 1^st^ cycle. p-values indicate a difference from 0 (one sample t-test). [E] Tone-evoked responses averaged over tones presented during 5-15 s (cycle#3 to cycle#6) [F] Source analysis results computed from the data in [E] The image is a group F map for the RND > REG thresholded at p = 0.05 (uncorrected).

An additional repeated measures ANOVA on response magnitude (mean power between 100-200ms post tone onset) with regularity (REG vs RND) and tone position in the 2 ^nd^ – 6^th^ cycles (i.e., from tone #11 to tone #60) as factors revealed a main effect of regularity only (F(1,21)=4.634, η2=.181, p=.043), with no effect of tone position (F(1,49)=1.063, η2=.048, p=.359) or interaction of the two factors (F(1,49)=.937, η2=.043, p=.599). Though clearly noisy, this tone-by-tone analysis reveals a sustained, stable difference between REG and RND conditions. As a control analysis, a repeated measures ANOVA on the first 10 tones in the sequence (cycle#1) indicated a main effect of tone position (F(1,21)=9.877, η2=.32, p<.001) only. Post hoc tests indicated that the responses to the first two tones are significantly different from the third through tenth tones (p<.01) in both REG and RND sequences, reflecting increased responses at sequence onset. Neither condition (F(1,9)=2.647, η2=.112, p=.119) nor the interaction of condition by tone position (F(1,9)=.556, η2=.026, p=.832) were significant. Together, these analyses confirm no difference between REG and RND during the first cycle (cycle#1), with a sustained difference between conditions emerging during the second cycle (cycle#2) onwards.

To further understand whether and how the tone-evoked responses in REG and RND contexts changed over time, we computed the mean evoked field differences between tones presented in the first and subsequent cycles in REG and RND conditions. Because responses to the initial couple of tones (first 2 tones in cycle#1) were affected by onset-response activity, we focused this analysis on the last eight tones of each cycle (cycle#1: tone 3-10; cycle#2: tone 13-20; and so on). The mean tone-evoked response (computed between 100-200 post onset) during cycle#1 was subtracted from that of cycle#2-#6 to understand how the presence of regularity affects tone responses. The data are plotted in Figure 5D. A repeated measures ANOVA with condition and cycle number as factors yielded a main effect of condition only (F(1,21)=4.723, η2=.184, p=.041). No effect of cycle number (F(4,84)=1.078, η2=.049, p=.373) or interaction of those two factors (F(4,84)=1.087, η2=.049, p=.368) was observed. This indicates a sustained difference between REG and RND conditions, that does not change over time. A one-sample t-test (uncorrected) confirmed that such differences for cycles#2-#6 in the REG condition were below zero, i.e. consistently *reduced* relative to cycle 1. [cycle#2 t(1,21)=-3.102,d=-.661,p=.003; cycle#3 t(1,21)=-3.288,d=-.701,p=.002; cycle#4 t(1,21)=-3.702,d=-.789,p<.001; cycle#5 t(1,21)=-2.161,d=-.461,p=.021; cycle#6 t(1,21)=- 2.478,d=-.528,p=.011]. In contrast, the same analysis for RND indicated non-significant effects [cycle#2 t(1,21)=-1.051,d=-.224,p=.153; cycle#3 t(1,21)=-1.7,d=-.363,p=.052; cycle#4 t(1,21)=-1.604,d=-.342,p=.062; cycle#5 t(1,21)=-1.829,d=-.390,p=.041; cycle#6 t(1,21)=- .125,d=-.027,p=.451].

Overall, the tone-evoked analysis demonstrates a consistent difference between tones presented in REG relative to RND contexts, the effect emerges early during the second regularity cycle (i.e. when the regularity has been established) and is manifested as a reduction in responses to REG tones, whilst responses to RND tones remain stable throughout the stimulus period.

Source localisation (see Figure 5F) for the contrast RND>REG during the tone-evoked response (full epoch – 0-200ms; extracted from the 3^rd^ cycle until sequence offset; 5-15 s; i.e. after the regularity in REG has been established; see Figure 4B and Figure 5E) identified sources in bilateral temporal lobe (superior temporal gyrus, Heschel’s gyrus) and bilateral IFG that underly the time-domain effect. The opposite contrast (REG>RND) yielded no significant activations.

**Table 1.**
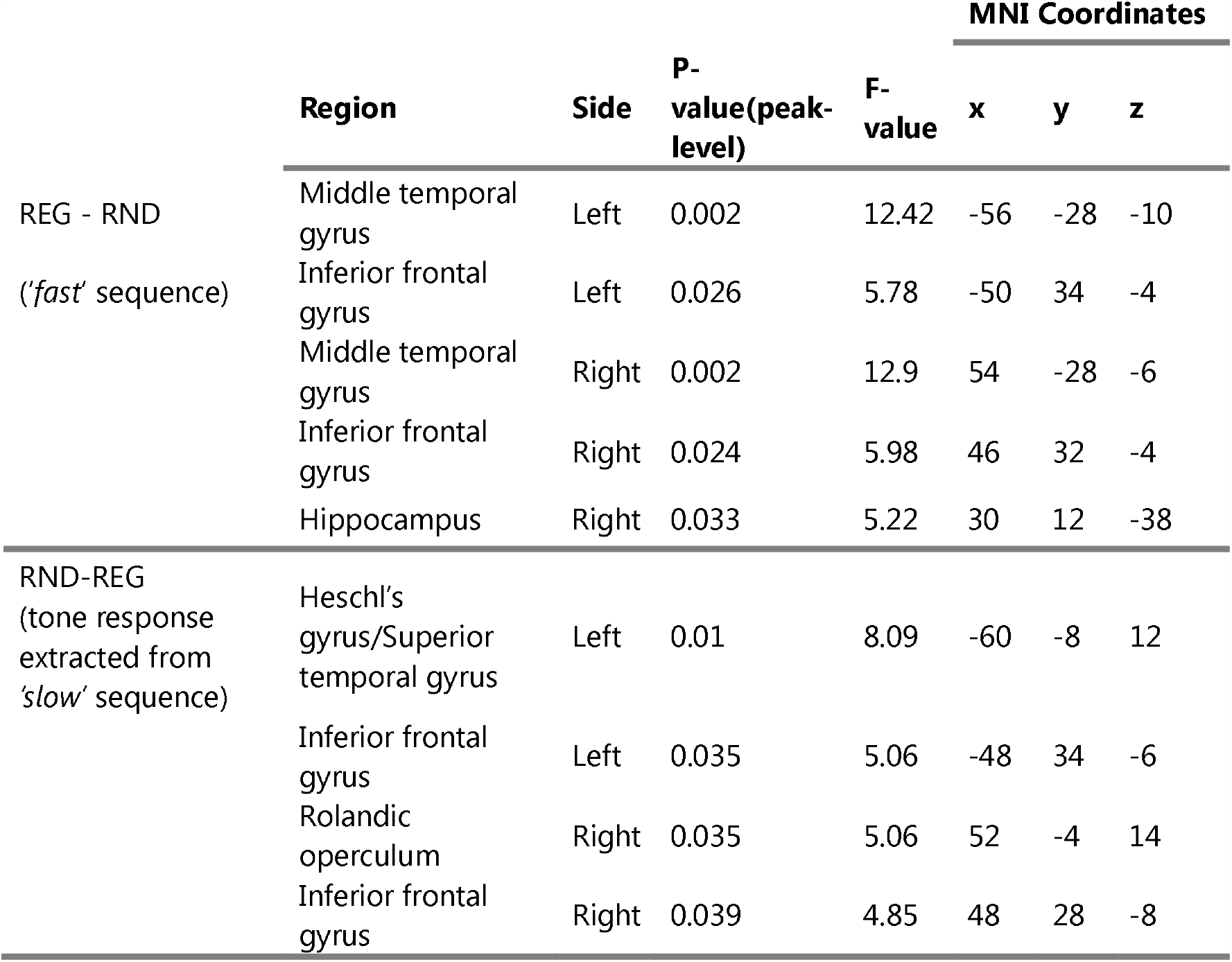
Summary of MEG source localisation results. MNI coordinates (x,y,z), and F values (p_voxel_ < 0.05). Anatomical labelling based on the Harvard-Oxford Cortical Structural Atlas.

### No significant correlation between tone-evoked and sustained-response effects

To investigate a potential link between the sustained response and tone evoked responses we correlated (spearman) the difference in the tone evoked response (REG-RND; mean power between 100-200ms post tone onset) with a difference in the sustained response (REG-RND; low pass filtered as in Figure 4B) during Cycle#2 and Cycle#6 across subjects. Both analyses yielded non-significant effects (p>0.2).

We also attempted more complex ridge regression analyses (Bates et al., 2015) over single trial data during Cycle#2 and Cycle#6 predicting the tone evoked response with the sustained response and trial number as predictors and subjects as random variable. No significant effects were observed (p>0.29).

## Discussion

We demonstrated that an increased sustained response to regular (REG) compared to random (RND) patterns previously observed in rapid tone sequences (20Hz; 500ms cycle duration), also occurs in slower sequences (4Hz; 2500ms cycle duration). This confirms the auditory brain’s remarkable implicit sensitivity to complex patterns. Critically, brain responses evoked by single tones exhibited the opposite effect - lower responses to tones in REG compared to RND sequences. The observation of opposing sustained and evoked response effects reveals parallel processes that shape the representation of unfolding auditory patterns.

### Sustained brain responses track pattern emergence even in slow sequences

Increased brain responses to predictable, relative to random patterns have previously been documented in many contexts (Barascud et al., 2016; Sohoglu and Chait, 2016; Southwell et al., 2017; Herrmann and Johnsrude, 2018; Herrmann et al., 2019). A greater amplitude for REG over RND stimuli is not easily interpretable as a response to physical attributes of the signal. Adaptation, for example, would result in the opposite pattern (Megela and Teyler, 1979; Pérez-González and Malmierca, 2014). Instead, the dynamics of this response, including when it diverges between REG and RND stimuli, suggest that the brain is sensitive to changes in the predictability of sound sequences. On an abstract level, observations regarding how the sustained response is modulated by sequence predictability suggest it might reflect the coding of precision, or *inferred reliability*, of the incoming sensory information (Barascud et al., 2016; Friston et al., 2017; Heilbron and Chait, 2018; Yon and Frith, 2021).

Here we showed that sustained response effects persist even when sequences are presented at a slower rate (4Hz). Despite the 5-fold increase in pattern duration, the divergence between REG and RND conditions occurred roughly at the same time (3 tones into the second cycle), in *slow* and *fast* sequences, consistent with ideal observer benchmarks (Pearce, 2005; Barascud et al., 2016; Harrison et al., 2020).

It is noteworthy that the sustained response was diminished in the *slow* compared to *fast* sequences. This could be attributed, at least in part, to limitations in human listeners’ memory capacity. Indeed, Barascud et al. (2016) observed a reduced sustained response to REG sequences consisting of cycles of 15 tones relative to 10 tones. This was interpreted as indicative of a threshold in encoding patterns that emerges when detecting longer repeating cycles. Similarly, Herrmann et al. (2019) reported reduced sustained responses in older individuals compared to younger participants, hypothesizing that this reduction could stem from age-related decline in tracking regularity patterns. To detect the emergence of regularity, the auditory system must presumably maintain and update a statistical model of the auditory input, registering tone repetitions, and decide at which point there is sufficient evidence to indicate a regular pattern. The efficiency of this process relies on the interplay between echoic and short-/long-term memory capacity (Bianco et al., 2020; Harrison et al., 2020).In our study, the introduction of gaps between consecutive tones and the subsequent increase in cycle duration from 500 ms to 2500 ms likely strained short-term memory capacity, leading to less precise memory encoding and therefore overall lower precision for the slow sequences. The behavioural results indeed indicate a decline in pattern detection (Figure 1). However, it is crucial to emphasize that despite this decline, the mean performance level remained high, underscoring the largely preserved sequence tracking capacity.

The brain mechanisms underlying the sustained response remain unclear. Source analysis suggests that the amplified response is driven by cortical activation in auditory, IFG and hippocampal sources (see also Barascud et al. (2016). A similar network involving the auditory cortex and IFG has been implicated in the generation of the Mismatch Negativity response (Näätänen et al., 2012) and has been postulated to represent the circuit responsible for maintaining an auditory model and conveying predictions to lower processing levels (Garrido et al., 2009; Heilbron and Chait, 2018).

According to one interpretation, the sustained response might reflect an excitatory processing mechanism, characterized by an increase in gain, potentially via neuromodulation, on units responsible for encoding reliable sensory information (Feldman and Friston, 2010; Auksztulewicz et al., 2017). In particular, tonic Acetylcholine (ACh) has been shown to be modulated by environmental uncertainty (Dalley et al., 2001; Yu and Dayan, 2005; Bland and schaefer, 2012). However, this interpretation may be less tenable, as it predicts heightened responses to tones within the REG sequences, which is contrary to our observed findings (see below). Alternatively, the sustained response may indicate an enhancement in the inhibition of neuronal units that convey low information content. This is consistent with prior research, albeit involving simpler stimuli, where an increase in inhibitory activity linked to the presence of predictable information has been documented (Natan et al., 2015, 2017; Schulz et al., 2021; Richter and Gjorgjieva, 2022; Yarden et al., 2022). Indirect evidence from dynamic causal modelling also suggests that the encoding of precision may be attributed to inhibitory mechanisms (Lecaignard et al., 2022). A specific role for inhibition, instead of excitation, in governing responses to predictable sensory stimuli, is also substantiated by behavioural findings: rather than capturing attention, predictable patterns are more easily ignored (Southwell et al., 2017) and are linked to reduced arousal (Milne et al., 2021). It is important to emphasize that M/EEG (or BOLD) do not readily differentiate between inhibitory and excitatory activity. Therefore, further advancement in understanding this phenomenon necessitates focused investigations at the cellular level.

### Reduced responses to tones in REG relative to RND patterns

Introducing temporal gaps between successive pips allowed us to disentangle the neural responses elicited by individual tones. Results revealed a reduction in neural activity in response to tones embedded within regularly repeating relatively to random patterns. This effect appears to be driven by relatively stable responses to tones in random patterns, but declining responses in the REG context. The dynamics of this effect are consistent with a step change in response magnitude during the second cycle (after the regularity has been introduced) that is then fixed for the remainder of the sequence.

Reduced response to REG tones is consistent with predictive coding theories (Rao and Ballard, 1999; Lee and Mumford, 2003; Friston, 2005, 2009). According to these models, top-down expectations, derived from statistical regularities in the external world, play a crucial role in suppressing anticipated sensory input. This mechanism serves as an efficient neural coding scheme, optimizing the allocation of neural resources and enabling the brain to prioritize the processing of novel or unexpected information, which may hold greater relevance (Olshausen and Field, 1996, 2004; Friston, 2005, 2009; Tang et al., 2018). Empirical support for these predictions, often referred to as ‘expectation suppression’, has been mounting across sensory modalities, (Baldeweg, 2006; Summerfield et al., 2008; Alink et al., 2010; Ouden et al., 2010; Todorovic et al., 2011; Kok et al., 2012; Todorovic and Lange, 2012; Barbosa and Kouider, 2018; Heilbron and Chait, 2018). In the auditory domain, Todorovic and de Lange (2012) demonstrated that when tones were expected based on the probability structure of tone transitions, they elicited suppressed auditory activity within a specific time window of 100–200 ms. This suppression was uniquely attributable to the phenomenon of expectation suppression and distinct from adaptation (repetition suppression) effects.

Notably, the effects we report manifest within this same time-window (120-200ms; during the M100 phase of the response). Whilst it is difficult to exclude low-level processes such as adaptation, several patterns in the dynamics of the development of these effects suggest that simple adaptation is unlikely to be a main factor. Firstly, the effects require processes that persist for 2500ms (duration of a cycle). Secondly, we do not see a gradual reduction in responses to REG tones that builds up over cycles. Rather there is a step change in the second cycle that is then consistent for the remainder of the sequence.

### Multiplexed representation of sequence predictability

The challenge faced by sensory systems is to accurately and swiftly represent information to support adaptive behaviour and facilitate interaction with the environment. A fundamental question pertains to whether the brain primarily encodes predictable or novel information (Press et al., 2020). Bayesian cognitive models propose that our predisposition to perceive what we expect enhances the fidelity of our sensory experiences (Wyart et al., 2012; Summerfield and de Lange, 2014; Kaiser et al., 2019). In contrast, cancellation models suggest that our perceptual system prioritizes unexpected stimuli, as they carry an informative value (Blakemore et al., 1998; Meyer and Olson, 2011; Richter et al., 2018). In line with these considerations, predictive coding models (Rao and Ballard, 1999; Friston, 2005, 2009) postulate the existence of two functionally distinct subpopulations of neurons within the brain. One encodes the conditional expectations of perceptual causes, while the other encodes prediction error.

Our findings confirm the coexistence of these facets of regularity coding within the MEG signal: the sustained response is consistent with the encoding of the predictability of the signal, whereas responses to individual tones appear to correspond to the coding of prediction error, as indicated by the reduced responses to predictable tones. Intriguingly, our results underscore the active involvement of the same neural network, encompassing the auditory cortex and the IFG, in both discovering structural patterns within auditory sequences and dampening responses to anticipated stimuli. However, the spatial resolution limitations inherent to MEG source analysis prevent definitive conclusions about the precise co-localization of these neural processes.

Indeed, the question of whether these manifestations stem from a singular process exhibiting differential characteristics in sustained and tone-evoked responses, or if they represent two distinct mechanisms, as proposed in previous works (Rao and Ballard, 1999; Friston, 2005, 2009), emerges as a crucial avenue for future exploration. For example, it is possible that the sustained response reflects activity linked to a tonic inhibitory drive (implementing gain control) onto sensory units, resulting in a diminished evoked response to individual stimuli. Notably, our study did not reveal a correlational relationship between tone-evoked and sustained responses. While this may tentatively suggest no direct linkage between the two mechanisms, it’s essential to consider the possibility that this observation could be influenced by the inherent noise in MEG measurements. We anticipate that more nuanced insights will be gleaned with the application of sensitive invasive tools in future investigations.

## Acknowledgements

We thank Theofilos Petsas for contributing to data collection. This work was supported by a BBSRC project grant to MC. RB. is funded by the European Union (MSCA 101064334). The funders had no role in study design, data collection and analysis, decision to publish or preparation of the manuscript.

